# Human Trophoblast Stem Cells Restrict Human Cytomegalovirus Replication

**DOI:** 10.1101/2023.12.13.571456

**Authors:** Tyler B. Rollman, Zachary W. Berkebile, Hiroaki Okae, Vivian J. Bardwell, Micah D. Gearhart, Craig J. Bierle

## Abstract

Placental infection plays a central role in the pathogenesis of congenital human cytomegalovirus (HCMV) infections and is a cause of fetal growth restriction and pregnancy loss. HCMV can replicate in some trophoblast cell types, but it remains unclear how the virus evades antiviral immunity in the placenta and how infection compromises placental development and function. Human trophoblast stem cells (TSCs) can be differentiated into extravillous trophoblasts (EVTs), syncytiotrophoblasts (STBs), and organoids, and this study assessed the utility of TSCs as a model of HCMV infection in the first trimester placenta. HCMV was found to non-productively infect TSCs, EVTs, and STBs. Immunofluorescence assays and flow cytometry experiments further revealed that infected TSCs frequently only express immediate early viral gene products. Similarly, RNA-sequencing found that viral gene expression in TSCs does not follow the kinetic patterns observed during lytic infection in fibroblasts. Canonical antiviral responses were largely not observed in HCMV-infected TSCs and TSC-derived trophoblasts. Rather, infection dysregulated factors involved in cell identity, differentiation, and WNT signaling. Thus, while HCMV does not replicate in TSCs, infection may perturb trophoblast differentiation in ways that could interfere with placental function.

**Importance:** Placental infection plays a central role in HCMV pathogenesis during pregnancy, but the species-specificity of HCMV and the limited availability and lifespan of primary trophoblasts have been persistent barriers to understanding how infection impacts this vital organ. Human TSCs represent a new approach to modeling viral infection early in placental development. This study reveals that TSCs, like other stem cell types, restrict HCMV replication. However, infection perturbs the expression of genes involved in differentiation and cell fate determination, pointing to a mechanism by which HCMV could cause placental injury.

## Introduction

Human cytomegalovirus (HCMV) is a highly prevalent beta-herpesvirus and a leading infectious cause of birth defects and adverse pregnancy outcomes. Recent surveys estimate the global seroprevalence of HCMV to be 83% across all demographics, and in the United States 65% of women of childbearing age are latently infected with HCMV (1). While HCMV infection rarely presents clinically in healthy individuals, the virus is an opportunistic pathogen that can cause severe illness in the immunocompromised and can infect the placenta and developing fetus. Congenital CMV (cCMV) infections occur in an estimated 0.66% of all live births in the United States and 10% of cCMV infections present as symptomatic at birth (2). Roughly 20% of children born with either symptomatic or asymptomatic cCMV develop sensorineural disabilities, such as neurosensory hearing loss, chorioretinitis leading to vision loss, microcephaly, and intellectual disability (3). Placental HCMV infections can cause a broad spectrum of injuries even in the absence of detectable viral transmission to the fetus. Pathologic findings in infected placentas include maldevelopment of chorionic villi, diffuse villitis, and areas of necrosis and calcification (4, 5). HCMV is a major contributor to idiopathic intrauterine growth restriction and an under recognized cause of fetal demise (6, 7). The timing of placental infection and virus-induced placental damage correlate with fetal injury (8–11). While placental infection can cause growth restriction or fetal demise after maternal infection at any time in pregnancy, the highest rates of HCMV-associated growth restriction and placental malformations are observed when infection occurs during the third trimester (12).

The placenta is formed by blastocyst-derived trophoblasts and these cells play a central role in antiviral immunity at the maternal-fetal interface. Trophoblast stem cells (TSCs) originate in the trophectoderm and small populations of TSCs are thought to be maintained throughout pregnancy. TSCs differentiate into proliferative cytotrophoblasts (CTBs), which are located both in cytotrophoblastic columns that anchor the placenta to the decidua and on the fetal side of chorionic villi. CTBs terminally-differentiate into extravillous trophoblasts (EVTs) and syncytiotrophoblasts (STBs). EVTs invade and remodel the maternal decidua, playing a key role in regulating maternal blood flow into the placenta. STBs form a non-proliferative, continuous syncytium that separates maternal blood from fetal circulation and release antiviral cytokines, type III interferons, and exosomes that contain antiviral microRNAs (miRNAs) (13). Type III interferon is constitutively expressed by trophoblasts during mid and late gestation, regulating the expression of select interferon stimulated genes (ISG) (14–17). The chromosome 19 miRNA Cluster (C19MC) encodes primate-specific miRNAs, which are expressed exclusively in the placenta, and have broad antiviral activity (14). These and other antiviral properties of trophoblasts make the placenta an effective barrier against the vertical transmission of most pathogens.

HCMV can infect and replicate in some trophoblast cell types despite their intrinsic immune defenses. In infected placenta, HCMV is most frequently detected by immunohistochemistry in CTBs and EVTs (5, 18). Experimental infection studies in explanted placenta and primary trophoblasts have revealed that HCMV can productively infect CTBs and EVTs but not STBs (8, 19, 20). How HCMV infection affects placental function remains unclear. Proliferative trophoblasts have an impaired capacity to differentiate when infected, which may interfere with the establishment and maintenance of the maternal-fetal interface and potentially facilitate viral dissemination from maternal circulation to the fetus (8, 19, 21). Inflammation triggered by viral infection can also cause placental dysfunction (22). In murine models of congenital Zika syndrome, fetal wastage may be driven by type I interferon signaling triggered by viral infection (23). Maternal polyinosinic:polycytidylic acid (Poly I:C) treatment, which triggers innate immune responses by stimulating pattern recognition receptors and directly activates intrinsic antiviral pathways such as protein kinase R, can affect placental and fetal development in mice and rats [reviewed in (24)]. Elevated concentrations of proinflammatory cytokines are observed in amniotic fluid and maternal blood in cases of cCMV (25–27). *In vitro* studies in tissue explants and organoids have revealed that HCMV infection triggers distinct patterns of cytokine production from decidua and placenta (28–30).

Several barriers have prevented a greater understanding of HCMV pathogenesis in the placenta. The cytomegaloviruses are species-specific and cannot replicate in animals other than their natural host. Tissue explants and primary trophoblasts are prepared from tissue that is collected either after elective pregnancy terminations or birth. While term placentas are typically discarded and can be collected for research purposes with relative ease, the use and collection of tissue from other times in gestation is more highly regulated if possible at all. Gene expression studies have focused on placentas aged up to 20 weeks or full-term; other data from the late second and third trimester comes from placentas that were delivered prematurely (31). Trophoblast cell lines such as BeWo and JEG3 have abnormal antiviral properties relative to primary trophoblasts that limit their utility as infection models (32, 33).

Methods to culture and maintain undifferentiated human trophoblast stem cells (TSCs) were recently developed (34). TSCs were first derived from blastocysts and villous tissue from first-trimester placentas, but cell lines were subsequently established from naïve embryonic or induced pluripotent stem cells and older placentas (30, 35). Mutant, clonal TSC lines can be generated by CRISPR/Cas9 mutagenesis and TSCs can be differentiated into EVT and STB monocultures or placental organoids (30, 34–37). Given the potential utility of TSCs as an experimental model of HCMV infection in the first trimester placenta, we investigated the capacity of HCMV to infect and replicate in TSCs and TSC-derived EVT and STB. While all three cell types can be infected by HCMV, very little, if any, virus was produced from infected trophoblasts. Further investigation revealed that HCMV replication was restricted at some point after entry. TSC growth was not affected by high multiplicity HCMV infections, and the virus was not maintained in the cells upon passage. While these findings may suggest that TSCs could function as an infection-resistant reservoir of proliferative cells, gene expression profiling did note significant changes in the transcription of factors related to cell fate and development, suggesting that infection may affect the capacity of TSCs to differentiate.

## Results

### HCMV infects but does not replicate in TSCs and TSC-derived trophoblasts

To assess whether HCMV could replicate in TSCs and TSC-derived trophoblasts, female (CT27) and male (CT29) TSCs were grown in media that either maintained the cells in an undifferentiated state (self-renewal media) or stimulated differentiation into EVTs or STBs (34, 36). The cells were infected with TB40/Ewt-mCherry (HCMV mCherry), a clinical-like strain that maintains the broad tropism of wild-type HCMV, at a multiplicity of infection (MOI) of 0.1 (38). HCMV-permissive retinal pigment epithelial cells (ARPE-19) were infected in parallel. mCherry-expressing cells were observed when all three cell types were infected (**Fig. 1A**). When cell-associated and extracellular virus was titered, HCMV replication peaked at 3 days post infection (dpi) in ARPE-19 cells (**Fig. 1B**). Titers from infected TSCs and STBs were near the limit of detection throughout the experiment. Peak titers from infected EVTs were higher, near 1X10^3^ plaque forming units (PFU)/ml. However, this concentration of virus could be detected at 1 dpi, likely before HCMV could have completed a single cycle of replication, and may be residual input virus.

**Figure 1.**
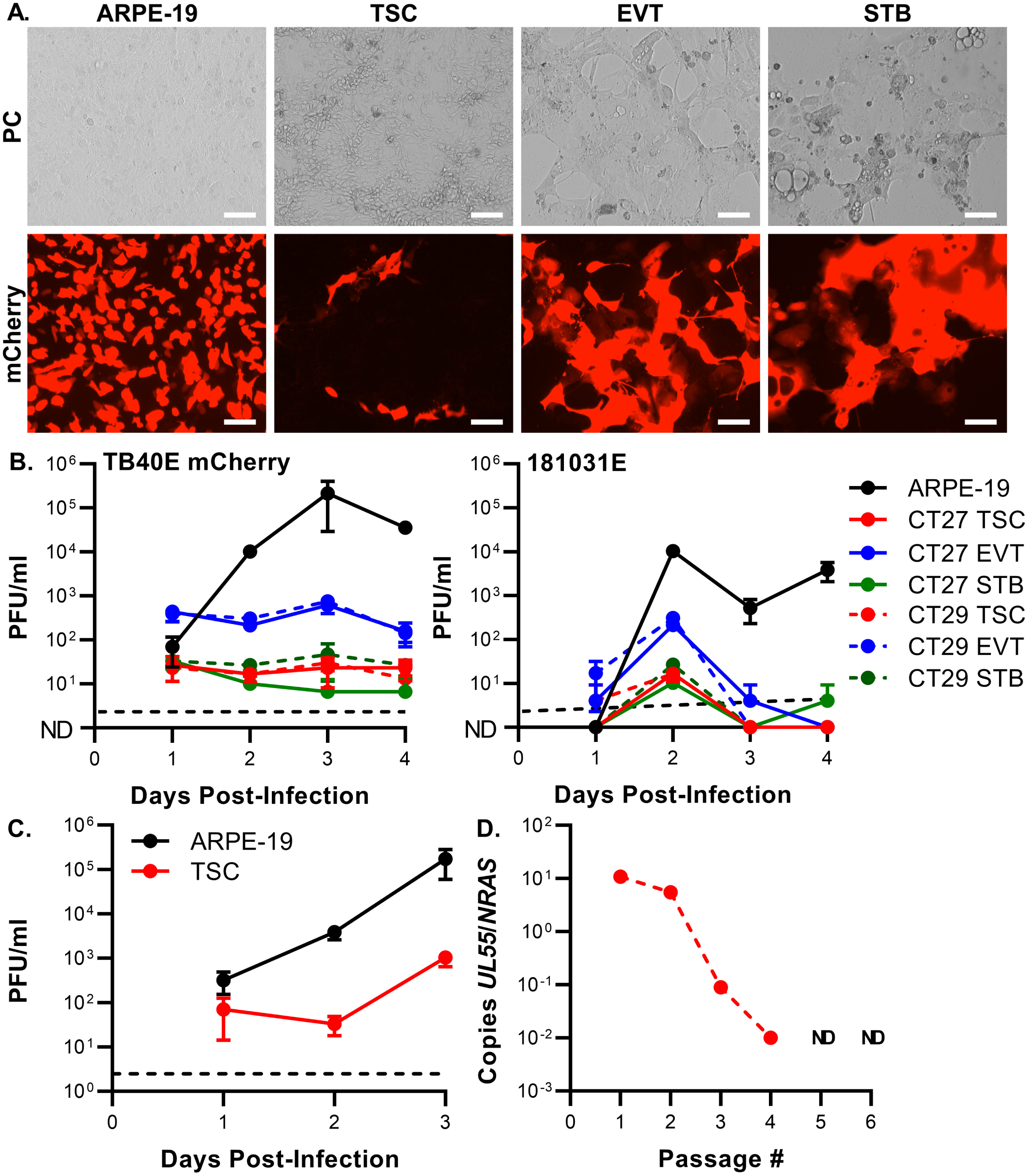
HCMV does not efficiently replicate in TSCs or TSC-derived trophoblasts. TSCs, TSC-derived EVTs and STBs, and ARPE-19 cells were infected with either HCMV mCherry or the HCMV clinical isolate 181031E at a MOI of 0.1. **(A)** Representative phase contrast and fluorescent micrographs show mCherry expression by HCMV mCherry-infected ARPE-19 cells and trophoblasts (CT29) at 2 dpi (Scale bar = 50 µm). **(B)** Cells were scraped and pooled with supernatant daily. HCMV production was measured by plaque assay on fibroblasts. (**C**) ARPE-19 cells were adapted to grow on TSC self-renewal medium for three passages and infected with HCMV. Virus production was measured as described above and compared with cells that had been grown under standard conditions. (**D**) HCMV mCherry-infected CT29s were passaged 1:10 to 1:15 six times. After each passage, the abundance of *UL55* and *NRAS* was quantified by droplet digital PCR. (ND: none detected).

HCMV mCherry is a TB40/E Bac4-based recombinant virus where non-excisable bacterial sequences and the *mCherry* expression cassette have replaced the viral genes *US1* through *US6* (39). To assess whether this gene replacement affects HCMV mCherry replication in trophoblasts, the cells were infected with the clinical isolate 181031E (**Fig. 1B**) (40). 181031E was isolated from the urine of a congenitally infected neonate and propagated exclusively on ARPE-19 cells to generate high-titer stocks after five passages. As with HCMV mCherry, very little 181031E replication was observed in trophoblasts relative to ARPE-19 cells. Taken together, the inability of HCMV to replicate in TSCs and TSC-derived cells does not appear to be a strain-specific defect.

TSC, ETV, and STB are propagated in media that includes several factors that are not routinely used when HCMV-permissive ARPE-19 cells and fibroblasts are cultured, including the histone deacetylase inhibitor valproic acid, the TGF-β inhibitor A83-01, and the Rho-associated protein kinase inhibitor Y27632. To test whether HCMV replication was inhibited by one or more of these factors, ARPE-19 cells were grown in TSC self-renewal media for 3 passages and infected with HCMV mCherry at an MOI of 0.1. While HCMV titers were lower in ARPE-19 cells grown in TSC media relative cells grown under standard conditions, the virus could still replicate (**Fig. 1C**). TSCs grow rapidly and must be passaged every three to four days even when infected with HCMV at a high multiplicity. Thus, TSC overgrowth constrained the duration of our infection studies. To assess whether HCMV would replicate in TSCs over longer time scales and if the viral genome was maintained when cells were passaged, TSCs were infected with HCMV mCherry at a MOI of 1 and passaged a total of six times, splitting 1:10 or 1:15 every three to four days. DNA was extracted from the cells after each passage and the abundance of host and viral genomes was quantified using droplet digital PCR (ddPCR) reactions specific to *NRAS* and HCMV *UL55* (41). The relative abundance of viral genomes exceeded that of host genome for the first two passages but rapidly decreased after subsequent passages; HCMV was undetectable and apparently cleared by the fifth passage (**Fig. 1D)**. Together, our observations suggest that TSCs and their derivatives do not support robust HCMV replication.

### TSCs restrict HCMV replication after entry

Given that mCherry expression was evidence that TSCs, EVTs, and STBs were being infected by HCMV (**Fig. 1A**), we next assessed whether other viral proteins were produced in infected trophoblasts. ARPE-19 cells and TSCs were infected with HCMV-mCherry at a MOI of 1 and immunostained with antibodies specific to the immediate early and late viral gene products IE1/2 and pp65 (**Fig. 2**). In ARPE-19 infected cells, high rates of IE1/2-positivity were detected at 24 hpi and virtually all infected cells expressed pp65 and mCherry by 72 hpi. The proportion of infected TSCs that expressed any virally-encoded protein was comparably lower; an estimated 10-20% of the cells expressed IE1/2. While the number of infected TSCs that expressed pp65 and mCherry increased as time progressed, at most 30% of IE1/2 expressing cells also expressed the other viral proteins.

**Figure 2.**
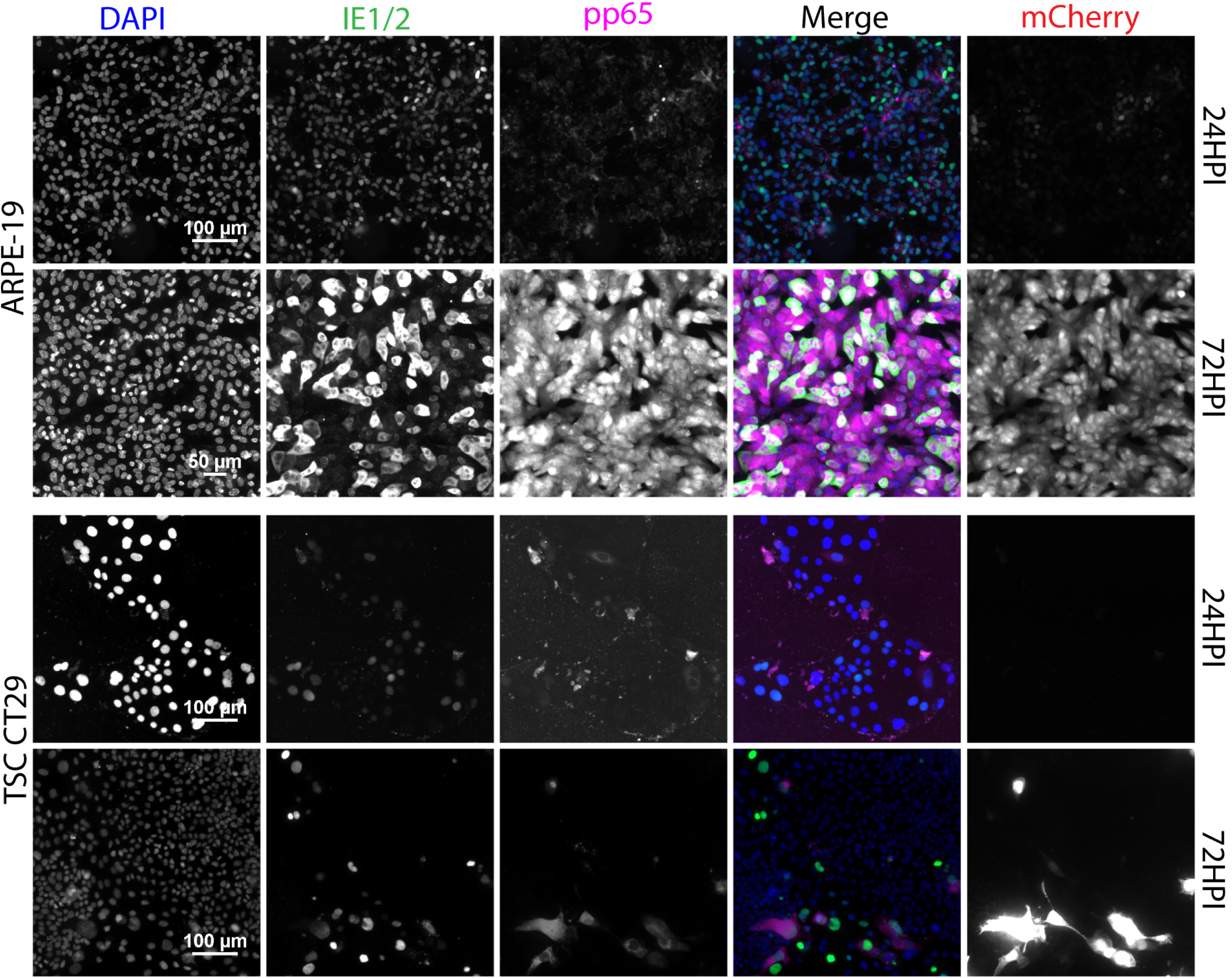
Fewer HCMV mCherry-infected TSCs express pp65 and mCherry than express IE1/2. ARPE-19 cells and TSCs were infected with HCMV mCherry at a MOI of 1 and immunostained with α-HCMV IE1/2 and pp65 at 24 and 72 hpi. Nuclei were stained with DAPI and the expression of virally-encoded mCherry was also assessed. The merged images show data from the DAPI, IE1/2, and pp65 channels and exclude mCherry data.

To better quantify the differences in viral protein expression between cell types, TSCs and ARPE-19 cells were infected with HCMV3F. HCMV3F encodes three fluorescent reporters with clearly distinguishable emission spectra that are linked by 2A peptides to HCMV proteins with immediate early (mNeonGreen-P2A-IE1/IE2), early (mTagBFP2-P2A-UL112/113), and late (mCherry-T2A-UL48A [small capsid protein]) expression kinetics (42, 43). TSCs and ARPE-19 cells were infected at low and high infectious doses (MOIs of approximately 0.05 and 0.5). Cells were analyzed by flow cytometry at 12, 24, 48, and 96 hpi. To measure viral protein expression over time, we first quantified mNeonGreen fluorescence in single cells. Using this reporter of immediate early protein expression to identify infected cells, mTagBFP2 and mCherry fluorescence was measured in mNeonGreen^+^ cells to determine whether infected cells also expressed early and late genes (**Fig. S1**).

In ARPE-19 cells, the abundance of mNeonGreen^+^ cells correlated with the initial infectious dose and increased over time (**Fig. 3**). At later time points, this reflects viral spread through the monolayer. The proportion of mNeonGreen^+^ cells that also expressed mTagBFP2 and/or mCherry also increased as time progressed (**Fig. 4**). Consistent with the results IE1/2 immunostaining, the proportion of TSCs that expressed mNeonGreen was 10-20% of the number of ARPE-19 cells that expressed the immediate early reporter at either infectious dose. The number of infected, mNeonGreen-expressing TSCs that also expressed either mTagBFP2 or mCherry was low even at late times post-infection. Thus, TSCs can be infected by HCMV, but these infections are largely unproductive and rarely progress past immediate early gene expression.

**Figure 3.**
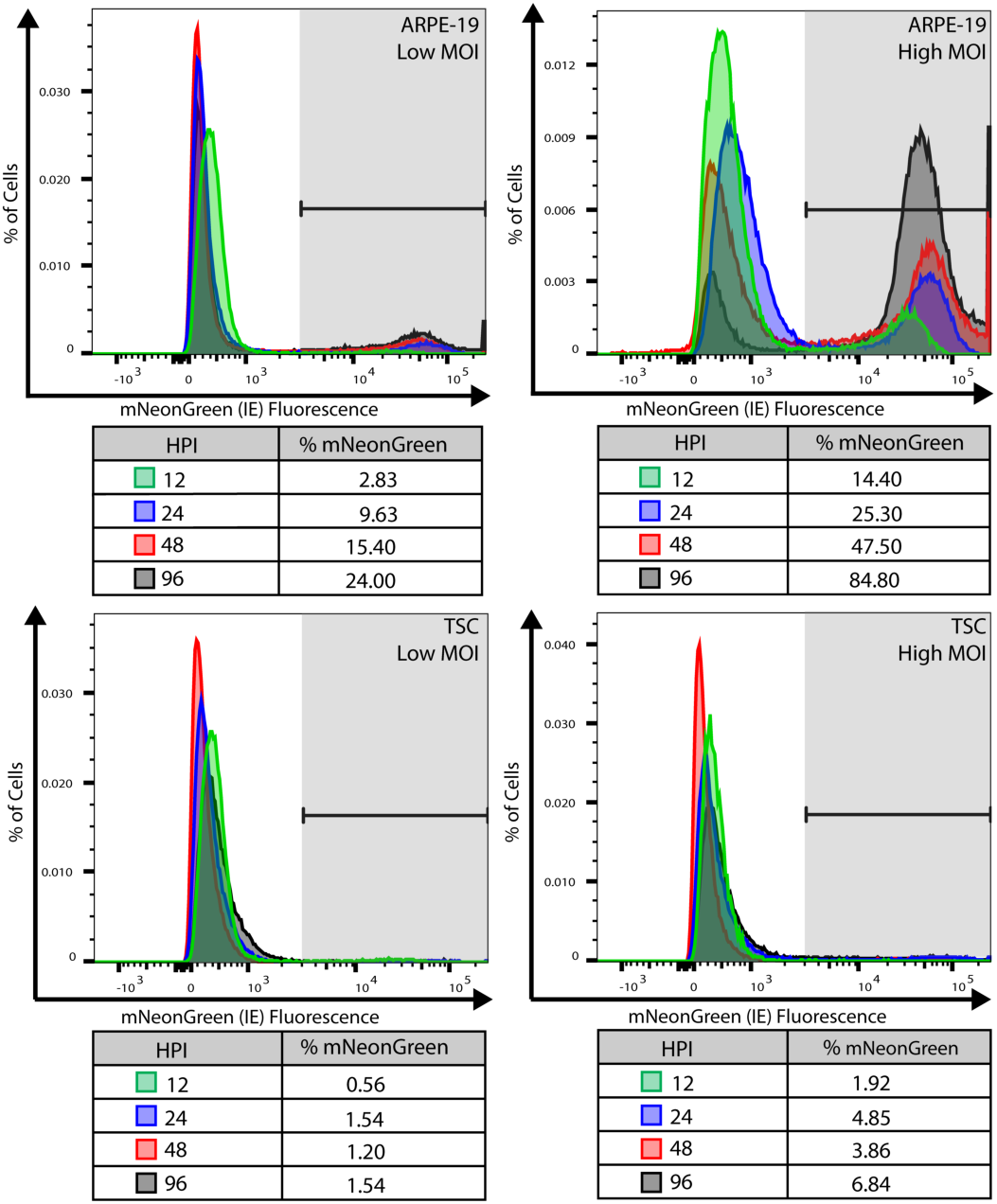
Fewer HCMV3F-infected TSCs express mNeonGreen than similarly-infected ARPE-19 cells. ARPE-19 cells and TSCs were infected at low and high MOIs with HCMV3F. Cells were harvested at 12, 24, 48, and 96 hpi and fluorescent protein expression measured by flow cytometry. mNeonGreen fluorescence is shown in the histograms, and the percent mNeonGreen-expressing cells in each population and time point is shown.

**Figure 4.**
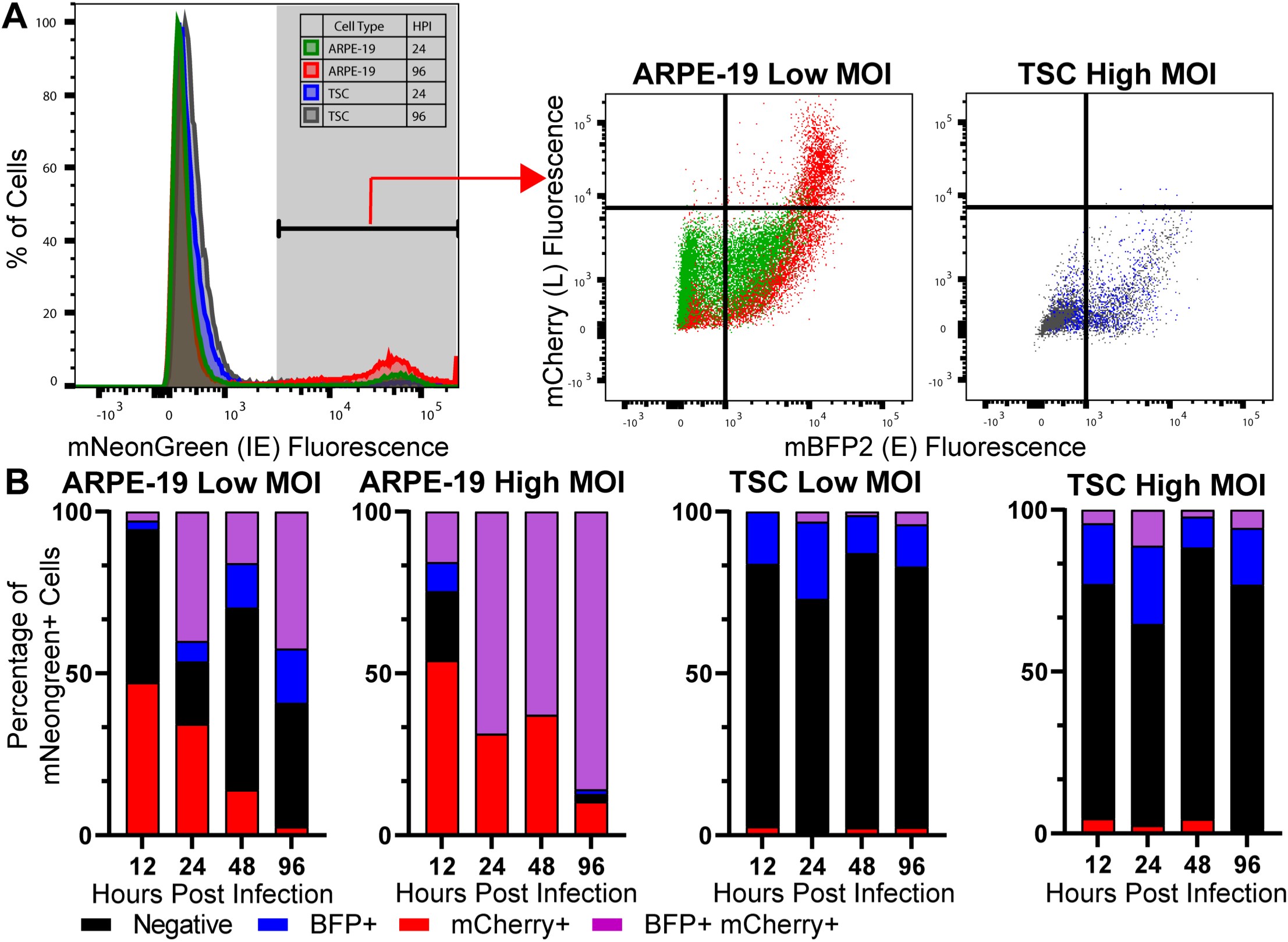
The expression of reporter proteins with early and late expression kinetics are suppressed in HCMV3F-infected TSCs. ARPE-19 cells and TSCs were infected with HCMV3F and the expression of virally-encoded fluorescent proteins was measured by flow cytometry. **(A)** Gating strategy used to quantify mTagBFP2 and mCherry fluorescence in mNeonGreen+ cells. Representative data from ARPE-19 low MOI and TSC high MOI infections at 24 and 90 hpi is shown. **(B)** The proportion of mTagBFP2 and mCherry-expressing cells, as determined by quad gating, is shown over time.

### HCMV gene expression is dysregulated in TSCs and TSC-derived cells

RNA sequencing (RNA-Seq) was used to study host and viral gene expression in mock- and HCMV-infected TSC and TSC-derived trophoblasts (**Fig. 5A**). TSCs, EVTs, and STBs were mock- or HCMV mCherry-infected at a MOI of 1 and RNA was extracted for analysis at 24, 48, and 72 hpi. For TSCs, cells were dissociated and fluorescence activated cell sorting (FACS) was used to separate the infected cells into populations with low or high mCherry fluorescence prior to RNA extraction (**Fig. S2**). RNA was extracted from EVTs and STBs directly as the two cell types uniformly expressed mCherry after infection. Reads were aligned to a concatenated reference sequence made up of the human and HCMV genomes (GRCh38 and TB40e-Bac4 [NCBI EF999921.1]).

**Figure 5.**
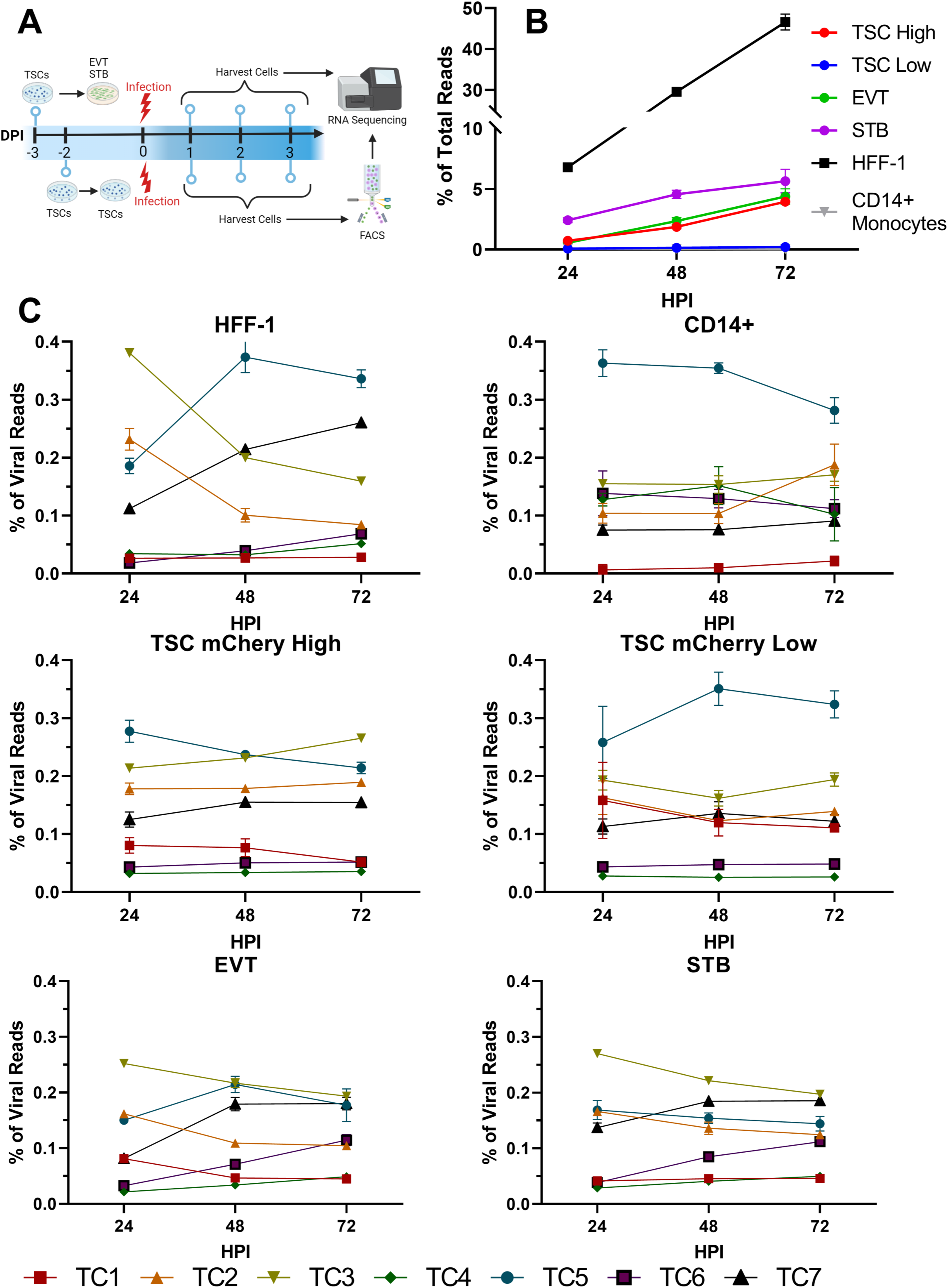
Infected TSCs and TSC-derived trophoblasts have distinct viral gene expression profiles when compared to infected fibroblasts or monocytes. TSC, EVT, and STB were infected with HCMV mCherry and RNA was extracted and sequenced at the indicated time points. Viral gene expression was compared with previously-published data from infected fibroblasts (lytic infection) and monocytes (latent expression) (44). (**A**) Summary of sample collection for RNA-sequencing. TSC-derived EVT and STB were grown in differentiation media for 3 days, mock- or HCMV mCherry-infected, and RNA was extracted at 1, 2, and 3 dpi. TSCs were infected similarly but sorted into populations with low or high expression of mCherry before RNA extraction. Figure created with the Biorender. (**B**) Illumina sequencing reads were mapped to the host and viral genomes. The proportion of reads that mapped to the viral genome at each time point is shown. (**C**) Viral reads were classified by temporal class and the abundance of each temporal class, expressed as a proportion of all viral reads, is shown over time.

The abundance of viral transcripts in infected trophoblasts was compared with RNA-seq data from a recent study of HCMV-infected fibroblasts (HFF1) and CD14^+^ monocytes, which model lytic and latent infections, respectively (44). First, the proportion of viral reads among all reads was compared across the five cell types (**Fig. 5B**). In infected fibroblasts, viral reads accounted for approximately 50% of all reads at 72 hpi. In contrast, very low levels of viral transcription was noted in latently infected monocytes and TSCs that expressed low levels of mCherry. TSCs that expressed high levels of mCherry and infected EVTs and STBs had an intermediate level of viral transcription, where the abundance of HCMV transcripts increased over time but peaked at a level that was 10-fold lower than was observed in fibroblasts.

Viral gene expression was next compared by temporal class (44). Most HCMV transcripts were assigned to one of seven temporal classes (TCs): immediate early (IE) genes (TC1), true early genes (TC2), delayed early genes (TC3), early-late genes (TC4), replication independent late genes (TC5), translation independent late genes (TC6), and true late genes (TC7). The proportional abundance of each TC was determined over time (**Fig. 5C**), and **Figure S 3** illustrates the expression of individual viral transcripts in a heat map. During lytic infection, IE transcripts were the most abundant before 24 hpi, early transcripts (TC2 and TC3) were the most abundant at 24 hpi, and late transcripts (TC5, TC6, and TC7) predominate at later times (44). Low-level expression of viral transcripts from all seven TCs was observed in latently-infected monocytes, where comparably little change in the relative abundance of the TCs occurred over time (44). Viral gene expression in infected EVTs and STBs was the most like lytic infection in fibroblasts. In these cell types, early transcripts (TC2 and TC3) were the most abundant at 24 hpi and late transcripts (TC6 and TC7) increased in abundance at 48 and 72 hpi. Two patterns of viral gene expression were noted in TSCs. There was no evidence of temporally coordinated viral gene expression in infected TSCs that expressed low levels of mCherry. In cells that expressed high levels of mCherry, high levels of early transcripts (TC2 and TC3) persisted until 72 hpi, when many late genes were not expressed to their expected level. Thus, RNA-Seq revealed an unusual pattern of viral gene expression in HCMV-infected TSCs, which is consistent with the defects in early and late protein expression that were observed by immunofluorescence and flow cytometry experiments.

### HCMV does not trigger a canonical antiviral response from infected trophoblasts

To assess how HCMV infection affects host transcription, gene expression in mock- and HCMV-infected cells was compared. Principal component analysis (PCA) revealed that the samples of TSCs and EVTs clustered according to when the cells had been collected (**Fig. 6A**). There was also separation between populations of mock- and HCMV-infected EVTs and STBs. Across all three cell types and time points, infection upregulated more transcripts than it downregulated (≥2fold, *p* < 0.05). Differential gene expression analysis revealed that HCMV infection had the greatest effect on transcription in STBs, where nearly 3000 genes were differentially regulated by 24 hpi (**Fig. 6B**). Infection had the least effect on TSCs that expressed low levels of mCherry; at most 394 transcripts were affected by HCMV in these cells. EVTs and TSCs that expressed high levels of mCherry had an intermediate phenotype: about 300 transcripts were differentially regulated in infected cells at 24 hpi and the number of differentially expressed transcripts increased at later times post-infection. Gene set enrichment analysis was used to illuminate host pathways that were affected by infection (45). The molecular functions and biological processes that were most highly enriched among infection-regulated transcripts were largely related to developmental processes, metabolic regulation, and cellular morphogenesis (**Fig. 6C**). Terms like “response to virus (GO:0009615)” that include transcripts that are canonically associated with an antiviral response were significantly enriched in some cases, but the absolute number of genes that were differentially regulated was low. Pathways related to inflammatory signaling and immune activation were enriched in some cell types and times.

**Figure 6.**
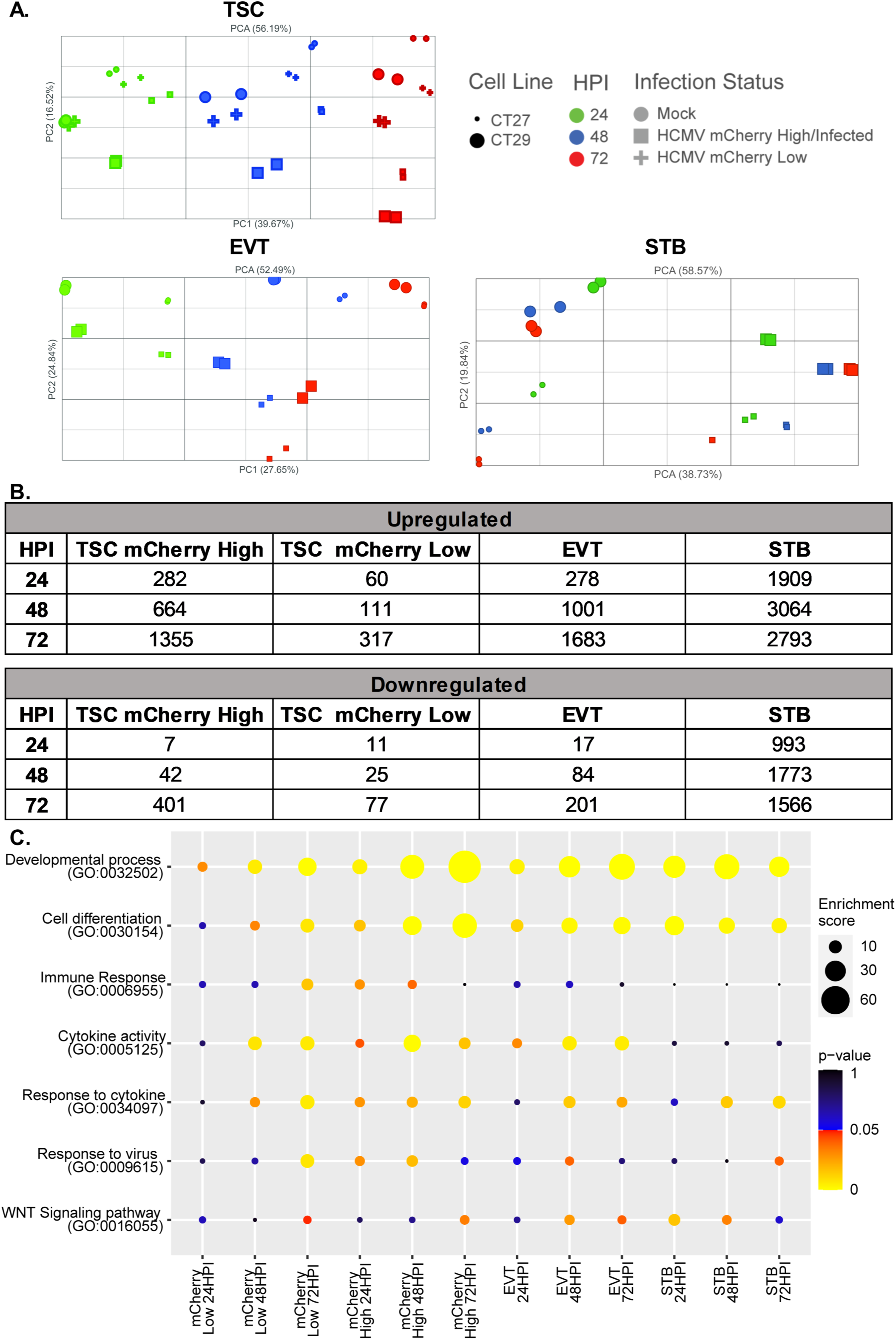
HCMV infection dysregulates more transcripts in differentiated trophoblasts than in TSCs. **(A)** PCA plots illustrate dimensionally-reduced gene expression in mock- and HCMV-infected TSCs, ETVs, and STBs. **(B)** Table summarizing the number of differentially regulated host transcripts in each cell type at 24, 48, and 72 hpi (≥2 fold, *p* < 0.05). (**C**) Gene set enrichment analysis was used to identify GO terms that were enriched among transcripts that were differentially expressed after infection. The results for select terms are illustrated by a bubble chart.

We next examined the expression of interferons and cytokines, interferon stimulated genes (ISGs), and factors involved in trophoblast differentiation or cell type specificity in infected and uninfected trophoblasts (**Fig. 7**). Primary trophoblasts constitutively express type III IFN, which helps to establish and maintain an antiviral environment in the placenta and decidua, while antiviral type I IFN responses have been found to cause placental dysfunction in some experimental systems (15, 23, 46). Type III IFN transcripts were detected at low levels in many samples, but HCMV infection did not significantly alter IFN expression. Gene ontogeny terms related to cytokine activity (e.g. GO:0005125, GO:0034097) were enriched in some infected trophoblasts. *IL1A, IL6,* and *IL11* were upregulated at every time point post-infection in EVTs and STBs. *IL11* transcription was also upregulated in TSCs that expressed high levels of mCherry, but this effect was only significant at 72 hpi. *TNF* was also significantly upregulated in these infected TSCs at 48 hpi.

**Figure 7.**
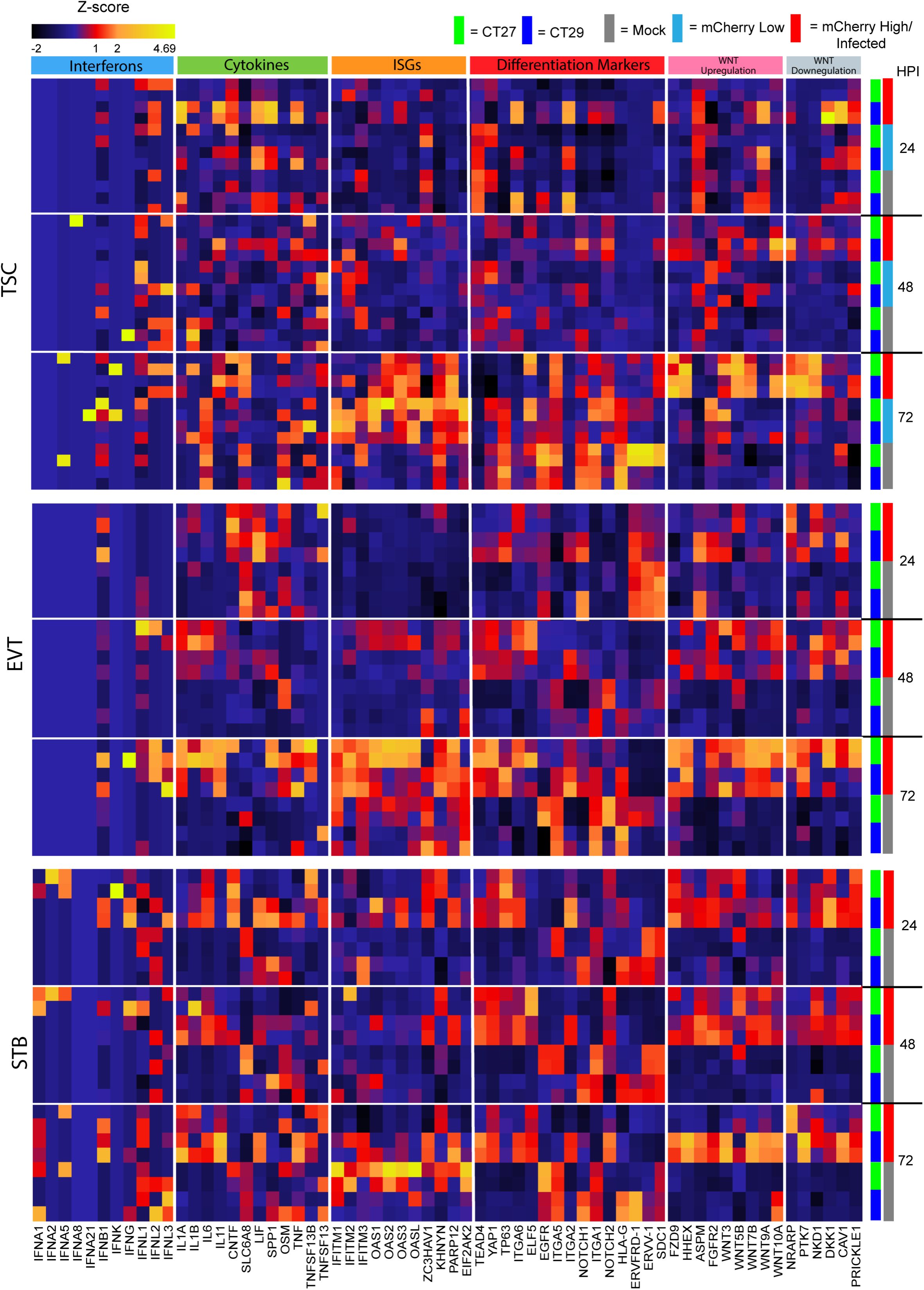
HCMV infection dysregulates transcripts involved in cell differentiation and inflammatory responses. Host gene expression was normalized using Z-scoring across all samples and organized by time point, infection status, and cell type. Representative transcripts encoding type I, II, and II interferons, IL-1 and IL-6 family cytokines, trophoblast biomarkers, and regulators of WNT signaling were assembled and grouped.

Interferons (IFNs) and pattern recognition receptors like RIG-I regulate the expression of hundreds of antiviral effectors, known as interferon stimulated genes (ISGs). Using an inclusive list that included 1905 different transcripts, we examined which ISGs were expressed by trophoblasts and how they were regulated by HCMV infection (47, 48). In TSCs, ISG expression was temporally regulated, and more transcripts were affected at late times post-infection and in cells that expressed high levels of viral mCherry. While some HCMV infection regulated ISGs encoded well known antiviral effectors like *IFIT5*, *MX1* and *OAS3,* many of the other transcripts have functions related to cell survival and proliferation. For example, *PRAME, NFIX,* and *A2M* are upregulated by infection and have known anti-apoptotic activities (49–51). *TRIB2* and *CCNA1* play key roles in stem cells self-renewal (52, 53). The greatest number of ISGs were differentially regulated in STBs (347, 519, and 505 at 24 hpi, 48 hpi, and 72 hpi respectively). Differentiated trophoblasts were like TSCs in that infection predominantly upregulated anti-apoptotic transcripts and factors that positively regulate cellular proliferation; *TRIB2* and *PRAME* were highly upregulated at all time points in infected STBs.

The expression of cell type-specific transcripts was analyzed to assess whether HCMV infection affected trophoblast differentiation. A recent analysis of differentiation trajectories in TSCs identified transcripts that could be used as markers of undifferentiated TSCs (*TEAD4*, *YAP1*, *TP63*, *ITGA6* and *ELF5*), differentiating EVTs (*ITGA5*, *ITGA2*, and *NOTCH1*), mature EVTs (*ITGA1*, *NOTCH2*, and *HLA-G*), and mature STBs (*ERVFRD-1*, *ERVV-1*, and *SDC1*) (**Fig. 7**) (54). All TSCs expressed some EVT and STB markers at 72 hpi. TSCs have been noted to express differentiation markers when grown in three-dimensional cultures, and the expression of these transcripts may reflect the spontaneous differentiation of some cells as cultures propagated in self-renewal media overgrow (54). HCMV infection significantly reduced the expression of *ERVFRD-1*, *ERVV-1, and HLA-G* in TSCs at 72 hpi. Biomarkers for both differentiating and mature cells were detected in all EVT samples and these transcripts were not differentially regulated by HCMV infection. However, transcripts associated with stemness, such as *ELF5* and *TEAD4,* were significantly upregulated in HCMV-infected EVTs. TSC markers were also upregulated in HCMV-infected STBs. Relatedly, Wingless/Integrated (WNT) signaling regulates the proliferation and maintenance of TSCs *in vitro*, and the removal of CHIR99021, a WNT activator, from self-renewal media triggers the differentiation of TSCs into EVTs (34, 54). Transcripts involved in WNT signaling pathways were among the transcripts that were most highly upregulated by HCMV infection. For example, *FZD9, NRARP, NKD1, WNT3, WNT5B,* were upregulated in infected TSCs. HCMV infection also perturbed WNT signaling in differentiated EVTs and STBs, where transcripts including *WNT3, WNT7B,* and *WNT9A* were upregulated. In summary, HCMV infection had more significant effects on host gene expression in differentiated EVTs and STBs than in TSCs. Infection did not appear to affect interferon transcription, but some inflammatory cytokines were upregulated by infection. While TSCs propagate after being infected with HCMV, transcripts involved in development and cell differentiation are dysregulated by infection; whether these changes in gene expression affect TSC function remains unclear.

## Discussion

TSCs are a recently developed and robustly characterized *in vitro* model of first-trimester human trophoblasts (34, 37). In this study, we used TSCs to model HCMV infection in the placenta, studying replication dynamics and the effects of infection on gene expression in undifferentiated and differentiated trophoblasts. We observed that TSCs and TSC-derived EVTs and STBs can be infected by HCMV but found little evidence of productive replication in any of the three cell types. High MOI HCMV infections have no apparent effect on the proliferation of TSCs and viral genomes are not maintained in the cells beyond the first several passages. Analyses of viral gene expression and protein production by immunofluorescence microscopy, flow cytometry, and RNA-Seq suggests that replication is inhibited after entry. Relative to lytic infection in fibroblasts, viral gene expression was notably dysregulated in TSCs. Infection also had muted effects on host transcription in TSCs when compared to similarly infected EVTs and STBs.

Our results were unexpected given decades of prior research that includes pathologic studies of HCMV-infected placentas and experimental infections done in tissue explants and primary trophoblasts. The broad consensus has been that HCMV infects and replicates in CTBs and EVTs but not STBs (5, 8, 18–20, 55–57). Chorion-derived trophoblast progenitor cells are likely the most similar cell type to TSCs that were previously used to study HCMV infection (58). HCMV replicates in these trophoblast progenitor cells, interfering with their ability to differentiate (19, 21). However, recent work has found that trophoblasts derived from the smooth chorion may be committed to a different cell fate than villous trophoblasts (59). Isolated reports have suggested that HCMV can replicate in STBs *in vitro* (60, 61). However, contaminating maternal and/or fetal epithelial cells can outgrow primary trophoblasts, and it is imperative to confirm that the intended cell type is being studied throughout an experiment when cells have been isolated from placenta (62). The CT27 and CT29 cell lines are advantageous in that the TSCs have been robustly characterized and meet many established criteria for first trimester trophoblast identity (34, 37, 63).

In this study, TSCs were infected with either a HCMV clinical isolate or one of two TB40E-based “clinical-like” laboratory strains. All viral stocks that were used were prepared by passaging virus a minimum number of times in epithelial (ARPE-19) cells and were confirmed to infect fibroblasts and epithelial cells with comparable efficiency. This was necessary because HCMV rapidly adapts during propagation *in vitro*, and commonly used laboratory strains (e.g. AD169 and Towne) and other viruses propagated extensively in fibroblasts can lack a glycoprotein complex that is required for HCMV’s naturally broad cell tropism (64). As the expression of virally encoded proteins confirmed that TSCs can be infected by HCMV, the cells likely either express one or more restriction factors that inhibit HCMV replication or are deficient in host dependency factors that the virus requires to replicate. HCMV can latently infect embryonic and induced pluripotent stem cells and can reactivate when the cells differentiate (65, 66). This suggests that HCMV may be generally unable to establish lytic infections in stem cells. TSCs may not be broadly restrictive to viruses as Zika virus can replicate in the cells (67).

Our findings are similar to observations made in HCMV-infected trophoblast organoids. Derived either directly from human placenta or from stem cells, trophoblast organoids are a three-dimensional model of the first trimester placenta (35, 68). Trophoblast organoids are largely refractory to HCMV infection but decidual organoids can be infected by the HCMV strains AD169r and TB40E (30). Like findings in two-dimensional TSC culture, trophoblast organoids can be readily infected by Zika virus (35, 67). Interestingly, ZIKV replicates less well in TSC-derived EVT and STB than it does in TSCs (67). Thus, the broad antiviral properties of primary trophoblasts may develop as the cells differentiate or as the placenta matures (15, 33, 69). Our gene expression profiling did not find strong evidence of type I or III interferon expression in TSCs nor or that these genes to be regulated by infection. Furthermore, while HCMV did upregulate some proinflammatory cytokines and ISGs in infected TSCs, EVTs, and STBs, the classical antiviral response to infection was more muted than is observed in other cell types.

HCMV infection early in gestation can cause placenta maldevelopment and early pregnancy loss and understanding how infection affects trophoblast differentiation has been an area of intense research focus (4). While HCMV infection did not effect the ability of TSCs to grow *in vitro*, we did not test whether infected cells could differentiate normally. Others have found that infected CTBs and chorionic villi have a diminished capacity to differentiate and invade matrigel (8, 21, 55). In trophoblast progenitor cells, proteins involved in cell cycle progression, pluripotency, and early differentiation are dysregulated after HCMV infection (19, 21). The HCMV-encoded cytokine cmvIL-10, an ortholog of human IL-10 that the virus expresses to manipulate the immune response to infection, impairs the invasiveness of infected trophoblasts (55, 70). WNT signaling also plays an important role in HCMV infection and trophoblast differentiation. HCMV sequesters and degrades beta-catenin in infected fibroblasts and SGHPL-4 cells, an EVT cell line, disrupting cellular function and potentially impairing EVT invasion during placental development (71). Notably, we found that *IL11*, which regulates trophoblast invasion, was upregulated by infection in most of the conditions we examined (72, 73). Given that RNA-Seq of TSCs, EVTs, and STBs revealed that the expression of cell type biomarkers is perturbed by HCMV infection and that WNT signaling is dysregulated, further research into how HCMV affects the capacity of TSCs to differentiate is merited.

## Materials and Methods

### Cells and viruses

ARPE-19 epithelial cells (CRL-2302) and HFF-1 human foreskin fibroblasts (SCRC-1041) were purchased from the ATCC (SCRC-1041) and propagated according to their instructions. CT27 and CT29 TSCs were maintained in an undifferentiated state and differentiated into EVT or STB as previously described (34). Briefly, cell culture plastics were coated with iMatrix-511 (ReproCELL MP892-011) recombinant laminin dissolved in TSC self-renewal media (0.05 µg/cm^2^) for at least 15 minutes before cells were seeded. TSC self-renewal medium contained DMEM/F12 (ThermoFisher 11320033) supplemented with 1% KnockOut Serum Replacement (ThermoFisher 10828010), 0.6% Penicillin-Streptomycin (ThermoFisher 15140122), 0.15% BSA (Millipore Sigma A1470), 1% Insulin-Transferrin-Selenium-Ethanolamine (ITS -X) (ThermoFisher 51500056), 0.2 mM L-ascorbic acid (Millipore Sigma, A8960), 25 ng/ml EGF (R&D Systems, 236EG200), 2 µM CHIR99021 (Tocris Bioscience 4423), 5 µM A83-01 (Tocris Bioscience 2939), 0.8 mM valproic acid (Millipore Sigma 676380), and 2.5 µM Y-27632 (Tocris Bioscience 1254). TSCs were dissociated by washing the cells with dPBS and incubating with a 1:1 mixture of dPBS and TrypLE Express Enzyme (ThermoFisher 12604013). TSCs were passaged every 2 to 3 days (upon reaching ∼80% confluence) at a ratio of 1:10 to 1:15.

For EVT and STB differentiations, cell culture vessels were coated with either 1 µg/ml (EVT) or 2.5 µg/ml (STB) of mouse Collagen IV (Corning™ 354233). For EVT differentiation, TSCs were seeded at a density of 7.8X10^3^ cells/cm^2^ in DMEM/F12 supplemented with 0.1 mM 2-mercaptoethanol (Gibco 21985-023), 0.5% Penicillin-Streptomycin (Gibco 15140-122), 0.3% BSA (Millipore Sigma A1470-25G), 1% ITS-X supplement (Gibco 51500-056), 100 ng/ml NRG1 (Cell Signaling Technology 5218SC), 7.5 µM A83-01 (Tocris Bioscience 2939/10), 2.5 µM Y27632 (Tocris Bioscience 1254/10), and 4% KnockOut Serum Replacement (ThermoFisher 10828-010), and 2% Matrigel (Corning 354234). At 3 days post-differentiation, EVT media was added that was formulated as described except in that it contained Matrigel at 0.5% and did not contain NRG. For STB differentiation, TSCs were seeded at a density of 2.1X10^4^ cells/cm^2^ in DMEM/F12 supplemented with 0.1 mM 2-mercaptoethanol, 0.5% penicillin-streptomycin, 0.3% BSA, 1% ITS-X supplement, 2.5 µM Y27632, 2 µM forskolin (Millipore Sigma F6886), and 4% KSR. STB media was replaced after 3 days.

TB40/Ewt-mCherry (HCMV mCherry) was a gift from Dr. Wade Bresnahan (University of Minnesota) (38). Infectious virus was reconstituted from the BAC by transfection into human fibroblasts by Dr. Bresnahan and passaged three times in ARPE-19 cells to generate the viral stocks used in this project. The clinical isolate 181031E was isolated from the urine of a congenitally infected child and passaged exclusively through ARPE-19 cells five times to generate the high titer stocks used in this project (41). The HCMV3F BAC was a gift from Dr. Luka Čičin-Šain (Helmholtz-Zentrum für Infektionsforschung) (42). Infectious virus was reconstituted by transfecting ARPE-19 cells with BAC DNA and Lipofectamine 3000 (ThermoFisher L3000001); high titer stocks generated after three passages in ARPE-19 cells were used for this project. The titers of HCMV stocks were calculated by TCID_50_ assays on both HFF-1 and ARPE-19 cells to ensure that broad tropism was maintained.

### Assaying HCMV infection and replication in trophoblasts

ARPE-19s, TSCs, EVTs, and STBs were seeded to 12-well plates as described above 3 days before infection. Each well was infected with 2.5X10^4^ PFU of HCMV for an approximate MOI of 0.1. Fluorescent micrographs of HCMV mCherry-infected cells were taken using an EVOS Cell Imaging System. At 1, 2, 3, and 4 dpi, cells were scraped into the media, collected, and flash frozen. To calculate HCMV titer, these samples were freeze-thawed a total of three times, serially diluted, and used to infect HFF-1 monolayers in plaque assays as previously described (74).

To assess the effects of TSC self-renewal media on HCMV replication, ARPE-19 cells were adapted to propagate in TSC self-renewal medium for 3 passages. These adapted and control cells were seeded onto a 12-well plate and infected with 2.5X10^4^ PFU of HCMV mCherry for an approximate MOI of 0.1. Cells were scraped into the media, collected, and flash frozen at 24, 48, and 72 hpi. To calculate HCMV titer, samples were freeze-thawed three times, serially diluted, and used to infect HFF-1 monolayers in plaque assays as previously described (74).

To assess the fate of HCMV-infected TSCs upon passage, TSCs were infected with HCMV mCherry at a MOI of 1. The cells were passaged every 3-4 days at a ratio of 1:10 to 1:15 a total of six times. After each passage, DNA was extracted from a portion of the remaining cells using the DNeasy Blood and Tissue kit (Qiagen). The relative abundance of human and viral genomes was assessed by droplet digital PCR using primers and probe specific to human HCMV *UL55* (FAM) and *NRAS* (Hex) as previously described (41, 75).

### Immunofluorescence assay

TSCs and ARPE-19 cells were seeded to an ibiTreat 4-well µ-Slide (ibidi 80426) at a density of 2X10^3^ cells/cm^2^ and grown overnight. The cells were infected with HCMV mCherry at a MOI of 1. After a 2 hour absorption, the cells were washed with dPBS and fresh media was added. Cells were immunostained at 24, 48, and 72 hpi. Media was aspirated and the cells were fixed with 4% paraformaldehyde (Sigma Aldrich P6148-1KG), permeabilized with 0.2% Triton X-100 (Sigma Aldrich 11332481001), and blocked with a 10% normal donkey serum (Sigma Millipore D9663-10). The cells were stained with mouse α-IE-1 (Ms. X pp72 mAb, Sigma 8B1.2) and rabbit α-pp65 (Biorbyt orb10511) primary antibodies at 1:200 dilutions, then washed with blocking solution. Donkey α-mouse AF488 secondary antibody (Jackson ImmunoResearch 715-545-150) and donkey α-rabbit Cy5 secondary antibody (Jackson ImmunoResearch 711-175-152) were used at a dilution of 1:250. After a final wash, the cells were mounted and nuclei stained using Mounting Medium with DAPI (ibidi 50011). Fluorescent micrographs were taken with a Nikon Ti-E Inverted Deconvolution Microscope and image analyses done with NIS-Elements Basic Research (version 4.60).

### Flow cytometry

Fluorescent protein expression vectors were cloned by PCR amplifying mNeonGreen (pCJB139 F: 5’ CACTAGTCCAGTGTGGTGGAATTGCCCTTACCATGGTGAGCAAGGGCGAG, R: 5’ TCGAGCGGCCGCCACTGTGCTGGACTTGTACAGCTCGTCCATGC) and mTagBFP2 (pCJB138 F: 5’ CACTAGTCCAGTGTGGTGGAATTGCCCTTACCATGGGACCTAAGAAGAAGAGAAAGG, R: 5’ TCGAGCGGCCGCCACTGTGCTGGAATTAAGCTTGTGCCCCAGTTTGC) from the HCMV3F BAC. The resulting amplicons were cloned into pKTS786 (a pcDNA 3.1/V5-His based vector) that had been linearized with BstX1 with an In-Fusion HD cloning kit (Takara 638909) (76). These vectors were transfected into ARPE-19 cells with Lipofectamine 3000 to use as single-stain controls for flow cytometry. ARPE-19 cells were infected with HCMV mCherry at a MOI of 0.1 and used as a single stain mCherry control.

TSCs and ARPE-19 cells were seeded at densities of 1X10^4^ cells/cm^2^ Twenty-four hours later, the cells were infected with HCMV3F at 1X10^3^ or 1X10^4^ PFU/cm^2^ for approximate MOIs of 0.05 and 0.5. After adsorbing for 2 hours, the infected cells were washed with dPBS and cell type-specific media added. Infected cells and single-stain controls were harvested by dissociation for flow cytometry at 12, 24, 48, 72, and 96 hpi. The dissociated cells were resuspended in a fixative buffer (0.5% BSA, 0.01% sodium azide in sterile dPBS). Cells were analyzed using a BD LSRFortessa X-20 Cell Analyzer with the same gating voltages for all time points. One hundred thousand events were recorded for each time point, and were gated in the following order: cells, single cells, single stain positive (for each fluorophore), and BFP+/mCherry+ cells within the mNeonGreen+ cell population. Flow analysis was done using FlowJo (version 10.8.0).

### RNA-Sequencing and bioinformatics analyses

For RNA-seq of HCMV-infected TSCs, 2X10^5^ CT27 or CT29 cells were seeded on T25 flasks and infected two days later with 7.5X10^5^ PFU of HCMV mCherry (approximate MOI of 1). Mock- and HCMV infected cells were trypsinized and collected at 1, 2, and 3 dpi, and stained with Ghost Dye Violet 450 (Tonbo Bioscences) for florescence-activated cell sorting with a FACSAria II cytometer (BD Biosciences). Cells were sorted directly into 2-Mercaptoethanol-containing RLT buffer and RNA was extracted using a RNeasy Mini kit (Qiagen). The cell populations used for RNA sequencing are summarized in **Table S1**. Sequencing libraries were prepared by the University of Minnesota Genomics Center using the SMARTer Pico Mammalian V2 kit (Takara). For RNA-Seq from EVTs and STBs, CT27 and CT29 were seeded into 6-well plates (7.5X10^4^ and 2X10^5^ cells/well, respectively) and differentiated for 3 days as described above. At 3 days post-differentiation, each well was infected at an approximate MOI of 1 with 2X10^5^ PFU of HCMV mCherry. RNA was isolated by directly lysing the cells with RLT buffer and using RNeasy Mini kits and used to prepare TruSeq Stranded mRNA (Illumina) sequencing libraries. Libraries were pooled and sequenced on an Illumina NovaSeq (S4 Flow Cell, 2×150-bp PE reads). A sequencing depth of >40M reads was obtained for TSC samples and >20M reads for EVT and STB samples; mean quality scores for all sequencing libraries were >Q30.

Sequencing data generated in this project was analyzed along with publically-available data from HCMV TB40E-GFP-infected fibroblasts and CD14+ monocytes (Gene Expression OmnibusGSE193467) (44). Sequencing reads were trimmed using Trimmomatic at default parameters (77) and aligned to a concatenated reference file that contained the human (GRCh38) and TB40e-BAC4 (NCBI EF999921.1) genomes with STAR version 2.7.2a using the default parameters (78, 79). Gene count tables were prepared with featureCounts (80). Gene expression analysis of host transcripts was done using PartekFlow (version 11.0.23.1004). After filtering out transcripts with fewer than 20 gene counts, transcript abundance was normalized using the median ratio method, and DESeq2 was used to identify differentially regulated transcripts when mock- and HCMV-infected samples were compared on a time and cell-type basis (81). Principal component analysis was used to dimensionally reduce and visualize data from each cell type. Differentially expressed genes were found to be statistically significant when they were upregulated or downregulated 2-fold or more, and DESeq2 calculated p-values were 0.05 or less. The expression of select genes of interest was visualized by heat map. To analyze the pattern of HCMV gene expression, viral reads were filtered from the host. The proportional abundance of each transcript in each sample was calculated by dividing its gene count by the total number of viral gene counts. RNA-Seq data have been deposited in NCBI’s Gene Expression Omnibus and are accessible through GEO Series accession number GSE248772 (82).

## Acknowledgments.

This project was supported by the Eunice Kennedy Shriver National Institute of Child Health and Human Development (R21HD087496 and R01HD109252 to CJB) and the Minnesota Masonic Charities (Masonic Cross-Departmental Grant in Children’s Health Research to CJB and VJB). TB40/Ewt-mCherry and HCMV3F were generously shared by Dr. Wade Bresnahan (University of Minnesota) and Luka Čičin-Šain (Helmholtz Centre for Infection Research). This work was supported by the resources and staff at the University of Minnesota University Imaging Centers (SCR_020997), the Genomics Center (https://genomics.umn.edu) and the Flow Cytometry Resource.

---

**Figure S1. Gating strategy used to analyze HCMV3F reporter protein expression**. Representative flow cytometry data is shown from the 24 hpi ARPE-19 Low MOI sample. Single cells were identified using forward and side scatter. After gating for mNeonGreen-expressing cells, the expression of virally encoded mBFP2 and mCherry was measured.

**Figure S2. Gating strategy for fluorescent activated cell sorting prior to TSC RNA-Seq.** TSCs were mock- or HCMV mCherry-infected. At 24, 48, and 72 hpi cells were dissociated and divided into populations that expressed high or low levels of mCherry using fluorescent activated cell sorting (FACS). Representative data from mock- and HCMV-infected TSC (CT29) cells collected at 72 hpi is shown. Cells were identified by forward scatter area versus side scatter area. Single cells were identified by using forward and side scatter heights first then secondarily gating by forward and side scatter widths. Single cells were gated by mCherry expression on the X-axis and Ghost Violet 450 on the Y-axis to sort live and dead cells, well as mCherry expressing cells. Three populations of cells were collected for RNA extraction and sequencing at each time point (summarized in Table S1): mock-infected mCherry^(-)^ cells, infected mCherry^(high)^ cells, and pools of infected mCherry^(-)^ and mCherry^(low)^ cells.

**Figure S3. Heat map illustrating normalized HCMV gene expression.** The relative expression of HCMV gene products was normalized by dividing the abundance of the transcripts by the total number of viral reads in that sample and scaled using a standard Z-scale across all genes. Viral transcripts were organized by temporal class, cell type, and time post-infection. Similarly processed, previously published data from HCMV-infected HFF-1 and CD14+ Monocytes is shown to illustrate gene expression patterns during lytic and latent infections (44).

