## Supplemental Materials for "Human Trophoblast Stem Cells Restrict Human Cytomegalovirus Replication"

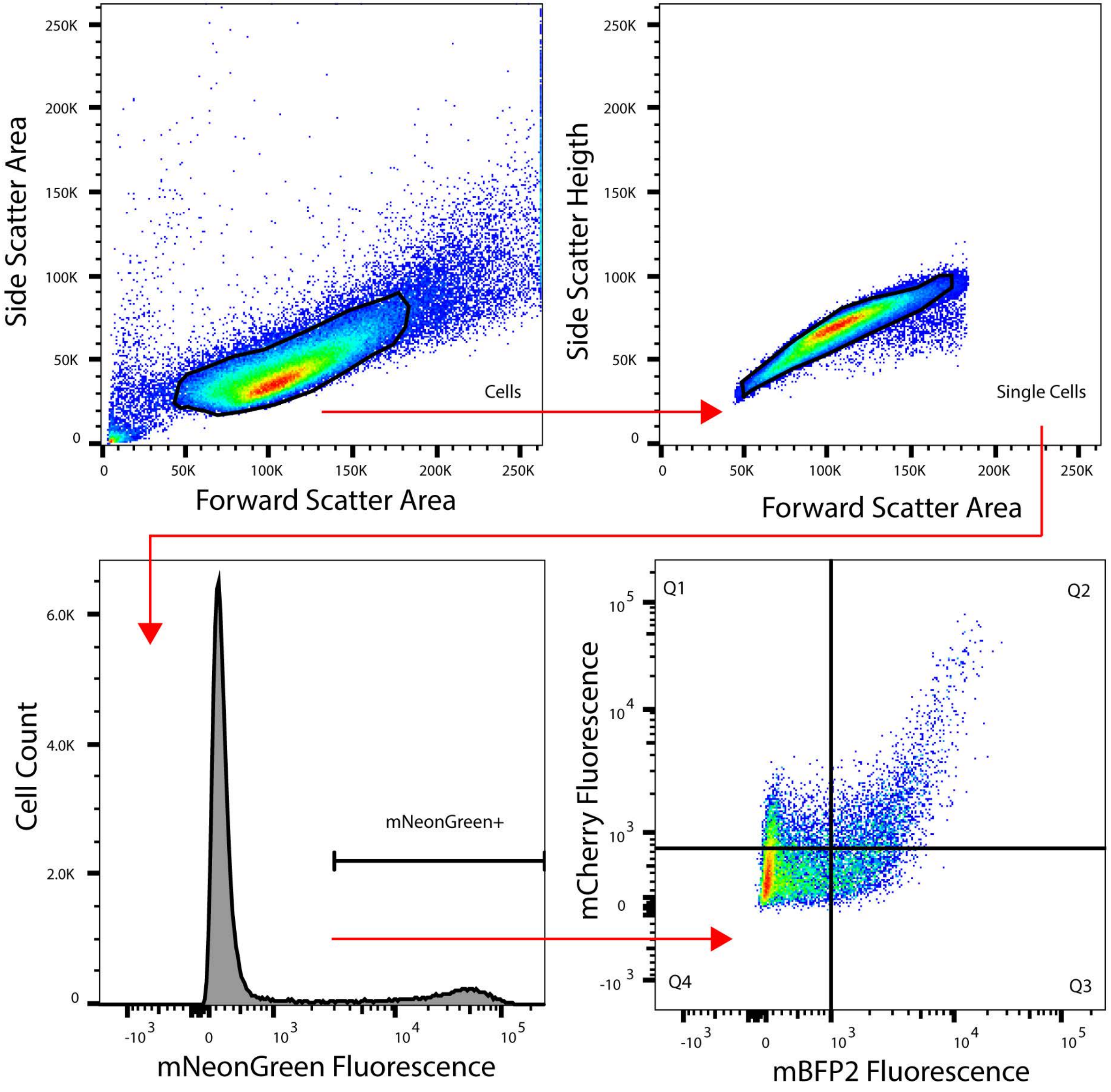
Side Scatter Area
250K
200K
150K
100K
50K
0
Cells
0
50K
100K
150K
200K
250K
Forward Scatter Area
Side Scatter Heigth
250K
200K
150K
100K
50K
0
Single Cells
0
50K
100K
150K
200K
250K
Forward Scatter Area
Cell Count
6.0K
4.0K
2.0K
0
mNeonGreen+
-10 3
0
10 3
10 4
10 5
mNeonGreen Fluorescence
mCherry Fluorescence
Q1
Q2
Q3
Q4
10 5
10 4
10 3
0
-10 3
-10 3
0
10 3
10 4
10 5
mBFP2 Fluorescence

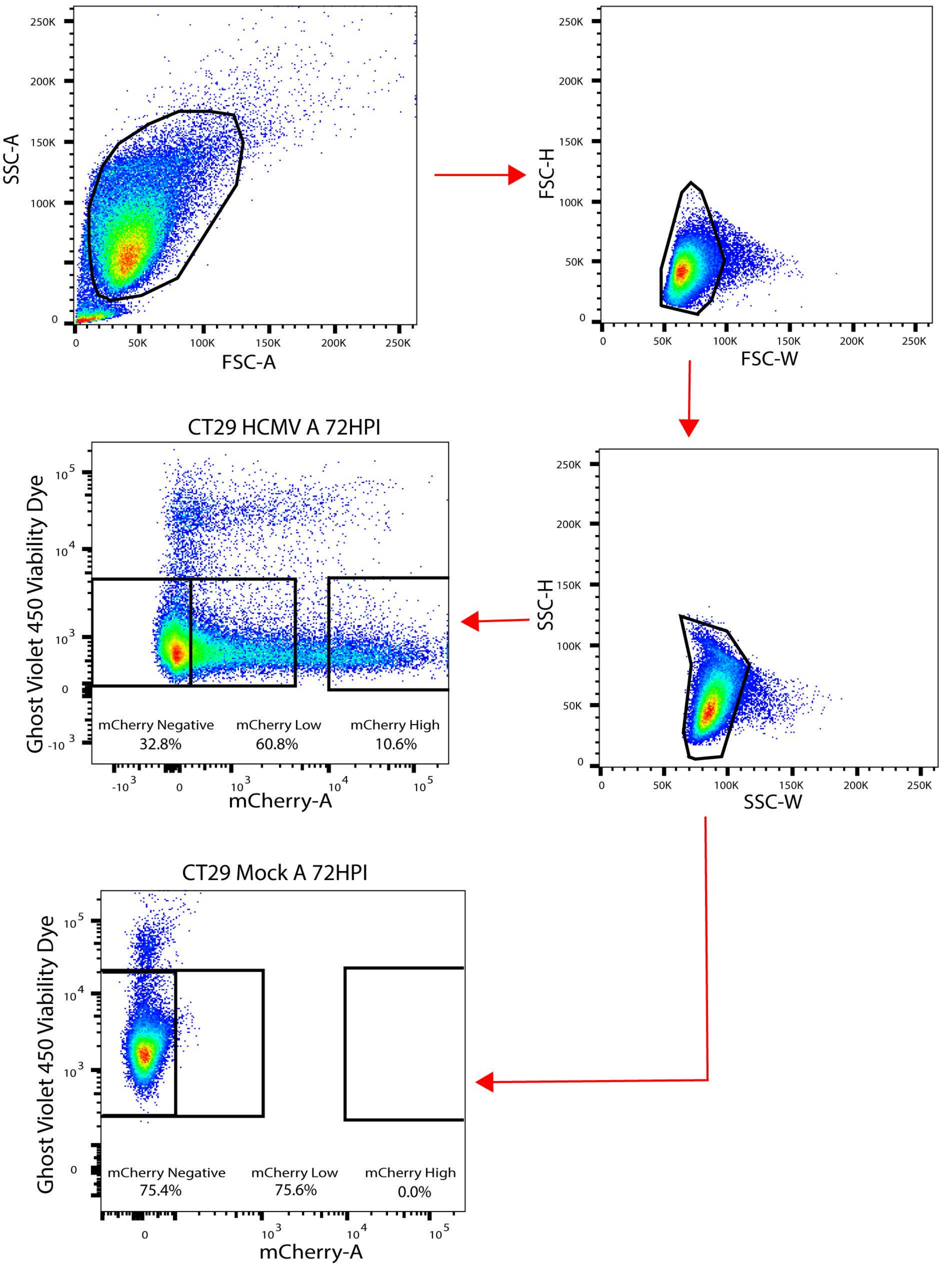
SSC-A
FSC-A
FSC-H
FSC-W
SSC-H
SSC-W
CT29 HCMV A 72HPI
Ghost Violet 450 Viability Dye
mCherry-A
mCherry Negative
32.8%
mCherry Low
60.8%
mCherry High
10.6%
CT29 Mock A 72HPI
Ghost Violet 450 Viability Dye
mCherry-A
mCherry Negative
75.4%
mCherry Low
75.6%
mCherry High
0.0%

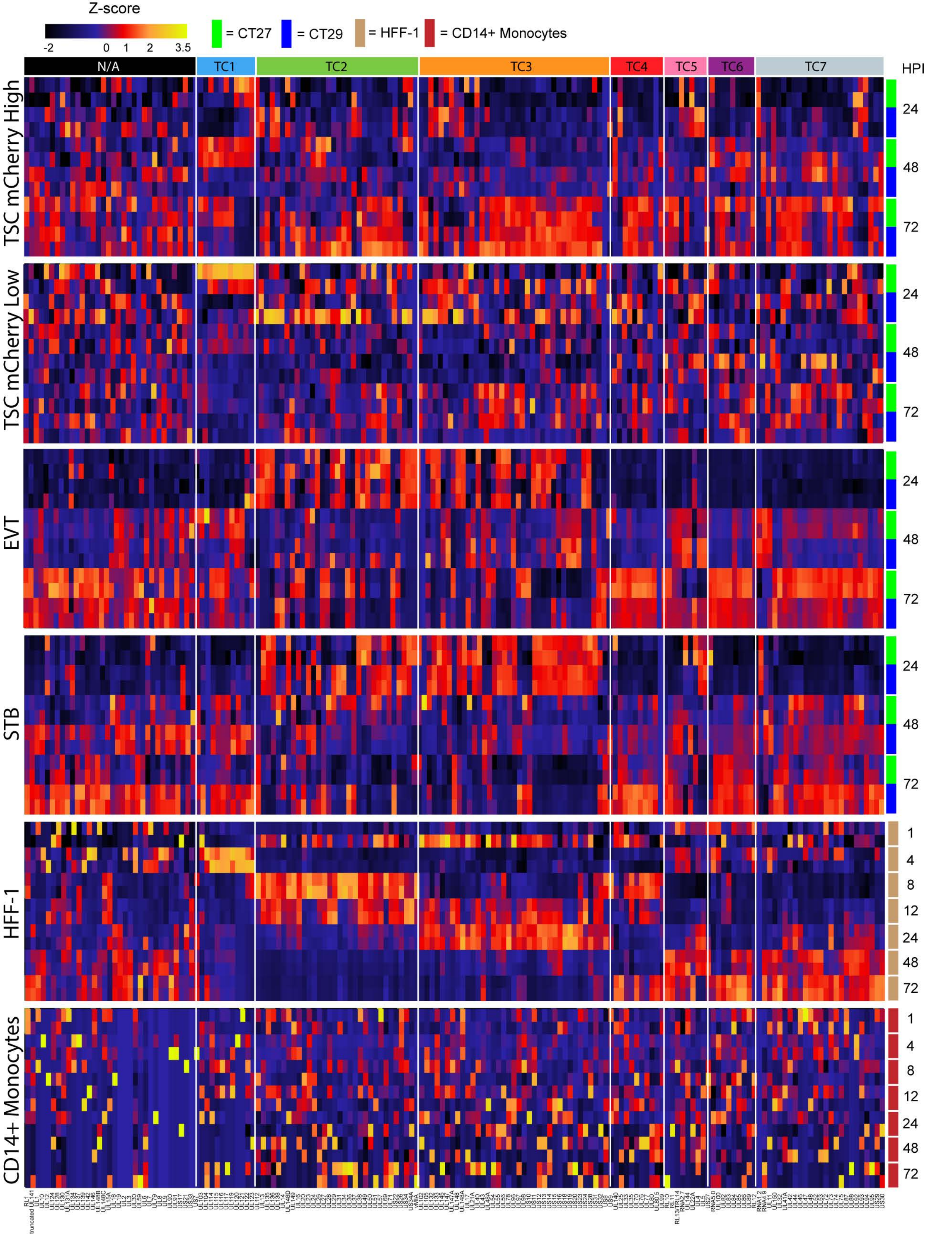
Z-score
-2
0
1
2
3.5
= CT27
= CT29
= HFF-1
= CD14+ Monocytes
N/A
TC1
TC2
TC3
TC4
TC5
TC6
TC7
HPI
TSC mCherry High
TSC mCherry Low
EVT
STB
HFF-1
CD14+ Monocytes
24
48
72
1
4
8
12

**Table S1. Infected TSC populations used for RNA-Sequencing**

| Cell Line | Infection | Replicate | Timepoint | mCherry (-) | mCherry Low* | mCherry High |
|---|---|---|---|---|---|---|
| CT27 | Mock | A | 24 | 72.8 | 77.0 | 0.0 |
| CT27 | Mock | B | 24 | 65.4 | 69.0 | 0.0 |
| CT29 | Mock | A | 24 | 81.0 | 84.5 | 0.0 |
| CT29 | Mock | B | 24 | 81.6 | 85.3 | 0.0 |
| CT27 | HCMV | A | 24 | 7.2 | 46.6 | 10.9 |
| CT27 | HCMV | B | 24 | 3.5 | 41.9 | 11.8 |
| CT29 | HCMV | A | 24 | 18.3 | 71.6 | 6.7 |
| CT29 | HCMV | B | 24 | 14.8 | 69.5 | 7.5 |
| CT27 | Mock | A | 48 | 77.9 | 79.9 | 0.0 |
| CT27 | Mock | B | 48 | 76.8 | 78.8 | 0.0 |
| CT29 | Mock | A | 48 | 83.1 | 83.5 | 0.0 |
| CT29 | Mock | B | 48 | 81.9 | 82.5 | 0.0 |
| CT27 | HCMV | A | 48 | 19.4 | 62.5 | 9.5 |
| CT27 | HCMV | B | 48 | 17.3 | 62.1 | 9.5 |
| CT29 | HCMV | A | 48 | 30.5 | 75.6 | 6.0 |
| CT29 | HCMV | B | 48 | 30.0 | 73.2 | 6.1 |
| CT27 | Mock | A | 72 | 70.2 | 70.5 | 0.0 |
| CT27 | Mock | B | 72 | 71.9 | 72.4 | 0.0 |
| CT29 | Mock | A | 72 | 75.4 | 75.6 | 0.0 |
| CT29 | Mock | B | 72 | 75.3 | 75.6 | 0.0 |
| CT27 | HCMV | A | 72 | 32.9 | 54.1 | 6.0 |
| CT27 | HCMV | B | 72 | 29.2 | 54.6 | 6.5 |
| CT29 | HCMV | A | 72 | 32.8 | 60.8 | 10.6 |
| CT29 | HCMV | B | 72 | 35.6 | 62.1 | 10.2 |

* The mCherry Low populations includes all mCherry (-) cells.

(grey box) Population used for RNA-Sequencing
